# Structure-based engineering of a nutrient acquisition protein enhances neutralizing antibodies and protection for the development of a gonococcal vaccine

**DOI:** 10.1101/2025.11.06.685839

**Authors:** Natalie Y.T. Au, En Ze L. Zhong, Jamie E. Fegan, Epshita A. Islam, Stacey X. Xu, Dixon Ng, Megha Shah, Laura-lee Caruso, Christine C.L. Lai, Anthony B. Schryvers, Scott D. Gray-Owen, Trevor F. Moraes

**Affiliations:** Department of Biochemistry, Temerty Faculty of Medicine, University of Toronto, Toronto, Canada; Department of Molecular Genetics, Temerty Faculty of Medicine, University of Toronto, Toronto, Canada; Department of Microbiology, Immunology, and Infectious Diseases, Cumming School of Medicine, University of Calgary, Calgary, Canada

**Keywords:** vaccine, TbpB, transferrin, antigen design, nutritional immunity

## Abstract

Gonorrhea is increasingly resistant to treatment and has been labelled an urgent threat due to the diminishing effectiveness of existing therapeutics. To address this challenge, we targeted the *Neisseria gonorrhoeae* transferrin binding protein B (TbpB), which is critical for iron acquisition and neisserial growth, as a vaccine target. Building on previous studies investigating the application of TbpB as an immunogen against various bacterial pathogens, we aimed to optimize this antigen for a broad protective effect. We compared the efficacy of wild type TbpB immunogens with engineered TbpB mutants that do not bind human transferrin (hTf) using infection studies in transgenic mice expressing hTf, which were required because the strict specificity of neisserial TbpB precludes its complexing with non-human transferrin. Comprehensive biophysical analyses confirmed that the introduced single residue mutations abolished hTf binding without compromising antigen structure. Immunization with the mutant antigens conferred increased resistance to infection by *N. gonorrhoeae* relative to that provided by the wild-type antigen in the humanized mice. When considering effector functions of the humoral response, we observed that the mutated antigen elicited more effective bactericidal and function-neutralizing activity. Through strategic mutations, we therefore enhanced vaccine effectiveness in a physiologically relevant model without significantly affecting the structure or immunogenicity of the antigen. This study highlights the use of rational structure-guided antigen design to drive effective immune responses and the potential interference of immunogen binding to host factors, and reinforces the utility of targeting TbpB in a gonococcal vaccine.

## 1. Introduction

Within the *Neisseriaceae* family, there are two known obligate human bacterial pathogens, *Neisseria meningitidis* (*Nme*) and *Neisseria gonorrhoeae* (*Ngo*)^1^. Significant reduction in the incidence of meningococcal-related meningitis and septicemia has been possible through the successful design and implementation of multiple vaccines^2–4^. In contrast, the prevalence of gonorrhea has steadily increased in recent years, with over 80 million new infections worldwide reported annually^5^. Paralleling the rapid increase in rate of infection is the emergence of multi-drug resistant strains, prompting this bacteria to be considered as an urgent public health threat^6,7^. Given the dwindling effectiveness of current therapeutics and the rise in antibiotic resistant strains, there is an urgent need to develop new prophylactic strategies^6^.

Numerous challenges continue to hinder the development of an effective vaccine against gonorrhea^8^. Natural infection does not induce long term immune responses, as reinfection with the homologous strain is possible, which prevents understanding of immune-protective mechanisms^9^. In addition, many conserved surface macromolecules, which are classically targeted for vaccine development, exhibit considerable variability due to antigenic and phase variation^6,9^. In recent years, highly conserved regions in prevalent molecules, like lipooligosaccharides (LOS), have been the target of vaccine design^10^. Alternatively, others have investigated outer membrane vesicle-based meningococcal vaccines due to the great similarities between the species^11^.

Currently, several clinical trials evaluating vaccine protection against *N. gonorrhoeae* are focused on repurposing a meningococcal B vaccine, 4CMenB (Bexsero), for gonorrhea, which has shown approximately 30-40% protection in retrospective studies^12–17^. While encouraging, a highly protective and cross-reactive vaccine is required to overcome the high diversity, adaptability and antimicrobial resistance the gonococcus has demonstrated over the past decades. To develop a more effective vaccine, antigen candidates should be highly conserved with an essential function and have the potential to elicit bactericidal immune responses.

Transferrin binding proteins (TbpA and TbpB) form a bipartite transferrin receptor located on the surface of all pathogenic *neisserial* strains, facilitating the uptake of the essential micronutrient iron from host transferrin^18^. The integral membrane protein, TbpA, functions as a gated pore required for extraction of iron from transferrin^18,19^, while the bilobed TbpB binds preferentially to iron-loaded glycoprotein transferrin to facilitate iron uptake^18–20^. The Tbp proteins are present in all strains of *Ngo* evaluated to date. In a strain lacking the lactoferrin receptor, which is variably expressed by some *Ngo* strains, the transferrin receptor is required for the pathogen’s colonization in a human male urethral challenge model^21^. Moreover, while TbpA is difficult to express, TbpB is a stable, soluble surface lipoprotein, making it an excellent candidate as a subunit vaccine^18^. Consistent with this, antisera raised against the Tbp proteins were bactericidal for *Nme* and conferred protection during a sepsis challenge^22^, and TbpB elicits a bactericidal response that protects against *Ngo*^23,24^.

Previous work to develop a TbpB-based vaccine targeting the porcine pathogen *Glaesserella parasuis* observed that a mutant variant that did not bind transferrin conferred protection that was superior to that observed by a wild-type TbpB and a commercial vaccine in a porcine challenge^25^. Such an effect is consistent with what has been seen with the *Nme-*derived factor H binding protein (FHbp), which is a component of both licensed serogroup B meningococcal vaccines, since reduced protection is apparent when mice express human factor H^26^. These results motivated us to test whether a binding-defective TbpB from *Ngo* would be a more effective immunogen than the wild type variant when animals express human Tf.

Leveraging structural data and predictive modelling tools, we engineered gonococcal mutant proteins that do not bind hTf in TbpBs from two gonococcal strains: MS11 and 48627^23^. These mutations preserved the structural integrity, as confirmed by comprehensive biophysical and immunological characterization of the mutants. To compare the protective efficacy of vaccines comprised of the wild type or binding-defective proteins, we employed transgenic mice expressing hTf, which allows potential interaction between the immunogen and its cognate host-derived protein^27^. While the antibody titres of these TbpB immunogens is indistinguishable, the antisera elicited by mutant TbpB variants is significantly better at blocking hTf binding, promoting complement-dependent killing and promoting opsonophagocytosis, and accelerates clearance of *Ngo* infection in the female lower genital tract. Collectively, our findings demonstrate the effectiveness of a rational structure-guided antigen design approach to develop improved vaccines with binding-defective proteins, and support the utility of using TbpB as a target to protect against *N. gonorrhoeae*.

## 2. Results

### Structural features of selected gonococcal TbpB

To guide the rational design of non-hTf binding gonococcal TbpB mutants, we selected the naturally-occurring sequence variants from strains MS11 and 48627, the former being a prototype strain commonly used in research and the latter a clinical isolate^23^. The TbpB variants from these share ∼94% amino acid similarity, and both represent the most prevalent cluster of TbpB variants identified by phylogenetic analysis, which were shown to induce a broadly cross-reactive antibody response across gonococcal strains^23^. Our attempts to crystallize the full-length proteins were unsuccessful, so we identified regions with high entropy using the Surface Entropy Reduction predictor (<u>SERp</u>) and then mutated these to reduce entropy without impacting functional surfaces in an attempt to aid our crystallization efforts^28^. The mutants generated allowed crystallization of the MS11 N-lobe (MS11^WT-Nlobe^) so that we could solve the structure at a resolution of 2.59 Å (Figure 1A, Table 1). Similar to other neisserial TbpB structures, the MS11^WT-Nlobe^ was comprised of an 8-stranded antiparallel β-barrel and a ‘handle’ domain that consists of 3 β-strands and a short α-helix (Supplemental Figure 1). Leveraging AlphaFold3, we confirmed that the predicted model of full-length MS11 TbpB (MS11^WT^) had similar structural characteristics to other TbpBs, with the full-length protein having an RMSD of 0.33 Å upon alignment to the 342 residues of MS11^WT-Nlobe^ (Supplemental Figure 1)^29^. With the N-lobe structure, we validated the overall fold of the functional lobe of the AlphaFold3 models of MS11^WT^ and 48627^WT^, both of which were predicted with high confidence (overall pTM >0.8). Moreover, loops localized in the cap region of the N-lobe, which are implicated in hTf binding, were predicted with confidence (Figure 1B,C).

**Figure 1.**
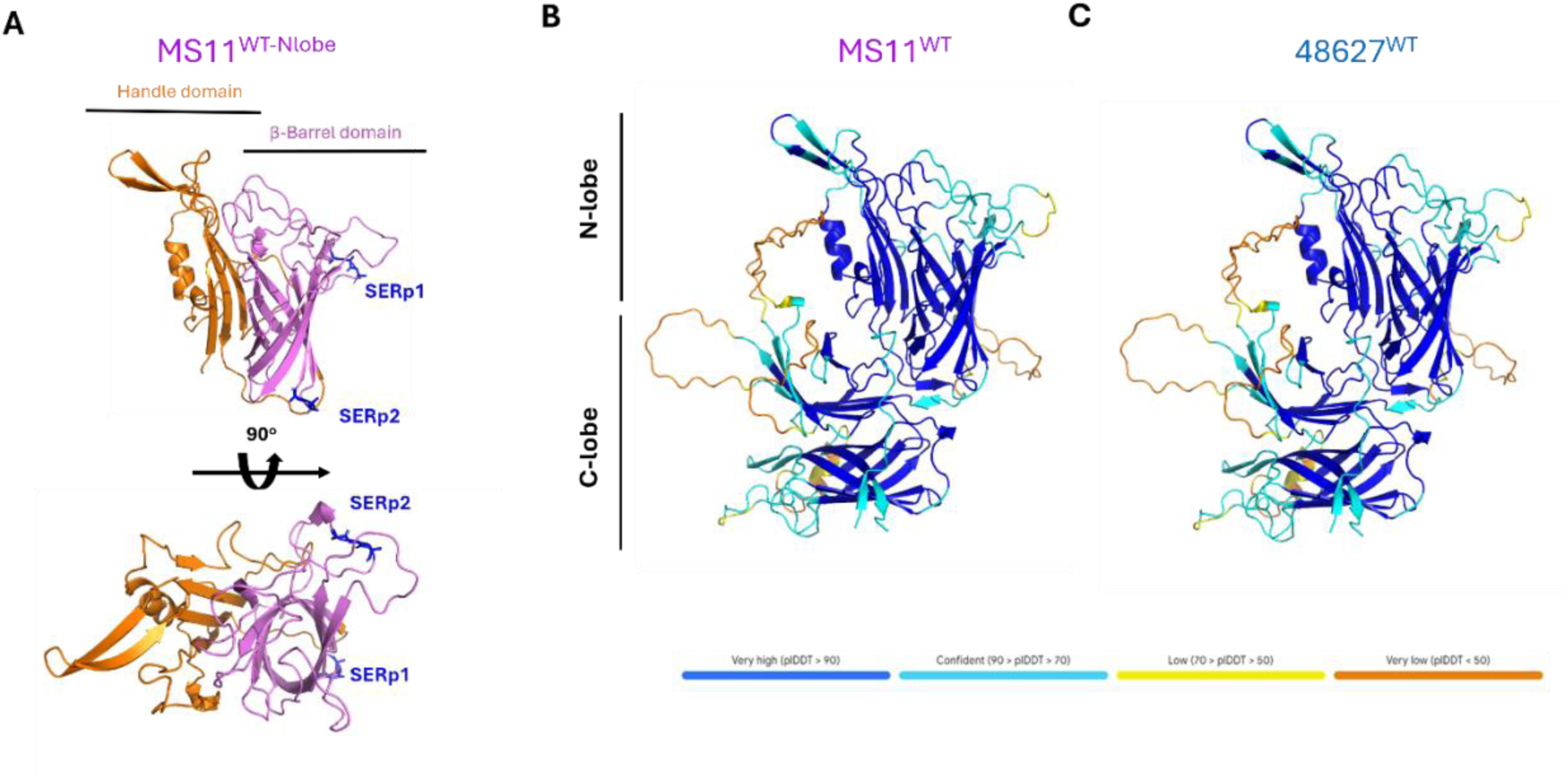
Structural models of gonococcal TbpBs. **(A)** X-ray crystal structure of the N-lobe from gonococcal MS11 TbpB. The N-lobe reveals an 8-stranded β-barrel domain (violet) and 3-stranded ‘handle’ domain (orange) connected through ten loops and turns. Clusters of residues with predicted high-entropy were mutated to alanine residues and are shown in blue as SERp1 and SERp2. Gonococcal MS11^WT^ **(B)** and 48627^WT^ **(C)** TbpB as predicted by AlphaFold3. Loops shown in yellow and orange were predicted with lower confidence (70>pIDDT), while regions in blue, such as the lobes, were predicted with higher confidence (pIDDT>70), as indicated by the legend below.

**Table 1.** Statistics of data collection and structural refinement for MS11^WT-Nlobe^ TbpB.

| **MS11<sup>WT-Nlobe</sup> TbpB** |  |
| --- | --- |
| *Phasing method* | MR |
| ***Data collection*** |  |
| *Space group* | C 1 2 1 |
| *a, b, c (Å)* | 202.668 60.182 183.284 |
| *α, β, γ (°)* | 90 110.71 90 |
| *Wavelength* | 0.9795 |
| *Resolution range* | 49.56 - 2.594 (2.63 - 2.59) |
| *Total reflections* | 63177 (2336) |
| *Multiplicity* | 3.3 (3.0) |
| *Completeness (%)* | 99.2 (91.9) |
| *Mean I/sigma(I)* | 5.62 (1.12) |
| *R-meas* | 0.17 (0.95) |
| *CC1/2* | 0.99 (0.63) |
| ***Refinements*** |  |
| *Reflections used in refinement* | 63177 (2336) |
| *R-work/R-free* | 0.1862 /0.1925 |
| *Number of non-hydrogen atoms* | 9770 |
| *Macromolecules* | 9266 |
| *Ligands* | 0 |
| *Solvent* | 504 |
| *Protein residues* | 1192 |
| *RMS(bonds)* | 0.003 |
| *RMS(angles)* | 0.65 |
| *Ramachandran favored (%)* | 94.92 |
| *Ramachandran outliers (%)* | 0.00 |
| *Rotamer outliers (%)* | 1.43 |
| *Average B-factor* | 45.32 |
| *Macromolecules* | 45.56 |
| *Ligands* | 0 |
| *Solvent* | 40.91 |

### Predicted models of TbpB variant structures implicate residues involved in binding function

While the crystal structure of MS11^WT-Nlobe^ provides insight on protein folding and supported the predictive model, the flexible surface loop structure precludes it from indicating which residues contact transferrin. As AlphaFold3 was not available at the time, we referenced the crystal structure of a meningococcal TbpB (from *Nme* strain M982) complexed (PDB: 3VE1) to hTf, previously solved by our lab^20^, to identify residues potentially implicated in transferrin binding(Supplemental Figure 2A). Although the meningococcal variant shares only 50% amino acid identity with the gonococcal TbpB N-lobe, functional loops have been shown to have some conserved interactions with hTf when comparing among the same serogroup of *Nme*^30^. Hence, by comparing the structures of this and our *Ngo* TbpB variants, we selected residues for mutagenesis in an attempt to generate binding-defective *Ngo* TbpB mutants.

The *Nme* TbpB interacts with the C-lobe of hTf through hydrogen bonds, salt bridges, and hydrophobic interactions. We therefore introduced mutations at homologous residues in *Ngo* TbpB to disrupt a predicted salt bridge, which we predicted would significantly reduce transferrin-binding affinity (Supplemental Figure 2B,C). Using a solid phase binding assay to assess hTf binding, we determined that a charge-reversal mutation at residue R200 in the MS11 variant corresponding to the meningococcal M982 R199, appeared to abrogate hTf binding (**MS11^R200E^**) (Figure 2A,B). Likewise, a mutation at the D209 residue in the 48627 TbpB (**48627^D209H^**), which corresponds to E222 on the M982 TbpB, caused a significant decrease in binding.

**Figure 2.**
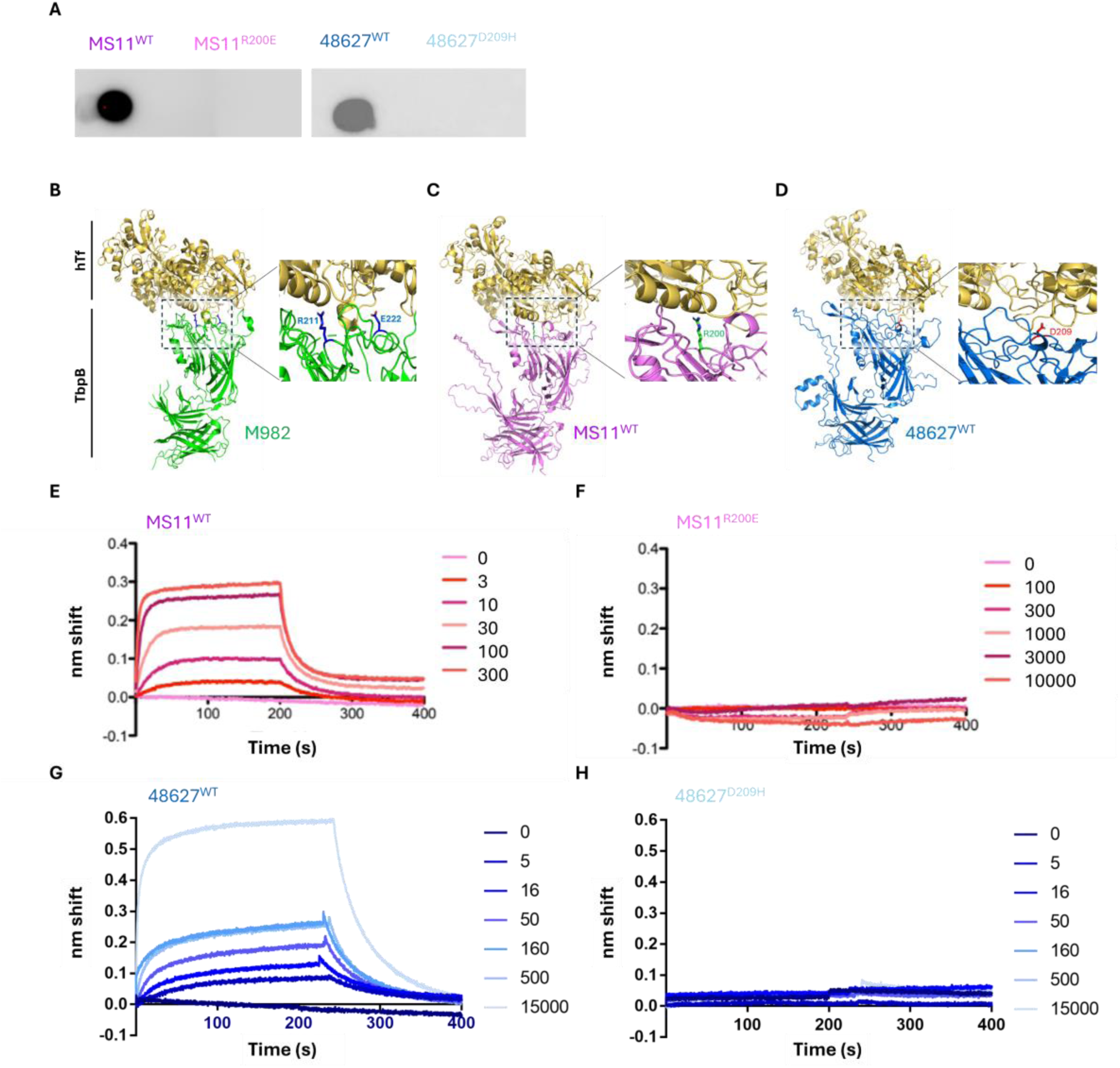
Design and validation of binding-defective gonococcal TbpB mutants. **(A)** Qualitative assessment of hTf binding for wild type and mutant TbpB variants through solid phase binding assay using hTf conjugated to HRP. **(B)** Co-crystal structure of meningococcal M982 TbpB complexed to human transferrin. M982 TbpB is shown in green and hTf is shown in yellow (PDB: 3VE1). The black box indicates the hTf binding interface, shown in the expanded illustration. Residues E222 and R211 on TbpB, involved in binding to hTf, are shown in blue. **(C)** MS11^WT^ TbpB -hTf complex as predicted by AlphaFold3. Residue R200 on TbpB is involved in hTf binding as shown in green. **(D)** 48627^WT^ TbpB – hTf complex models as predicted by AlphaFold3. Residue D209 is involved in binding to hTf as shown in red. Biolayer interferometry analysis of the binding of MS11^WT^ **(E)**, MS11^R200E^ **(F)**, 48627^WT^ **(G)** and 48627^D209H^ **(H)** TbpBs binding to iron-loaded (holo) hTf. TbpB was expressed with a N-terminal histidine tag and a biotin acceptor peptide (BAP) to aid in purification from *E. coli* lysate and for *in vivo* biotinylation, respectively. Concentrations of hTf (nM) tested are shown on the right, with an association of 200 seconds followed by dissociation in kinetics buffer (PBS with 0.1% BSA) for 200 seconds. Binding to hTf was assessed at concentrations up to 300 nM for the WT protein and up to 15000 nM for the mutant.

To expand upon our qualitative characterization of the hTf binding function of these mutants, we quantified binding affinity through biolayer interferometry^31^. The steady state affinity of the MS11^WT^ and 48627^WT^ TbpB variant for hTf was 18 ± 1.3 nM and 7.2 ±0.5 nM, respectively (Figure 2E,G) (Table 2). The point mutations that we introduced resulted in a dramatic decrease of binding affinity, as no binding to hTf was observed at µM concentrations for these mutants (Figure 2F,H).

**Table 2.** Kinetic analysis of TbpB-human transferrin binding data.

| **Replicate** | **MS11<sup>WT</sup>** | **48627<sup>WT</sup>** | **MS11<sup>R200E</sup>** | **48627<sup>D209H</sup>** |
| --- | --- | --- | --- | --- |
| 1 | 16.8 nM | 6.7 nM | n.d. | n.d. |
| 2 | 19.3 nM | 7.7 nM | n = 3 | n = 3 |
| 3 | 18.7 nM | 7.3 nM |  |  |
| Average | 18.3 ± 1.3 nM | 7.2 ± 0.5 nM |  |  |

\*Errors indicate standard deviation; n.d. = not determinable

### Mutation at the hTf binding interface does not affect the structural integrity of TbpB

Upon expression and purification of the mutant TbpB proteins, no differences in size were observed relative to their wild-type counterpart (Supplemental Figure 3). Our AlphaFold3 models did not predict any significant changes by the single residue changes that we made in each variant (Supplemental Figure 2), however it remains unclear whether this tool is sufficiently accurate to predict the impact of point mutations, particularly with respect to protein folding and stability^32,33^. We thus aimed to experimentally confirm that the loss of binding did not significantly affect the structural integrity of the antigen. Between the 48627 and MS11 variants, we prioritized characterization of 48627 due to its ability to elicit a broadly cross-reactive immune response^23^ and the absence of existing structural characterization for this protein.

Using nanoscale differential scanning fluorimetry (nano-DSF), we examined the thermal stability of the 48627^WT^ and its corresponding mutant, and observed no significant differences in their thermal unfolding profile (Figure 3A). We next tested structural integrity through limited proteolysis, treating 48627^WT^ and 48627^D209H^ variants with chymotrypsin and trypsin. Upon digestion with each of these enzymes, we observed a similar degradation pattern for the two proteins over time (Figure 3B). Within 30 minutes, over half of the full-length protein had been degraded by both enzymes, with no significant difference between wild-type and mutant protein (Figure 3C, D). Together, these findings suggest that the charge reversal mutation used to abrogate hTf binding for the 48627 TbpB variant does not significantly affect the structural integrity of the protein.

**Figure 3.**
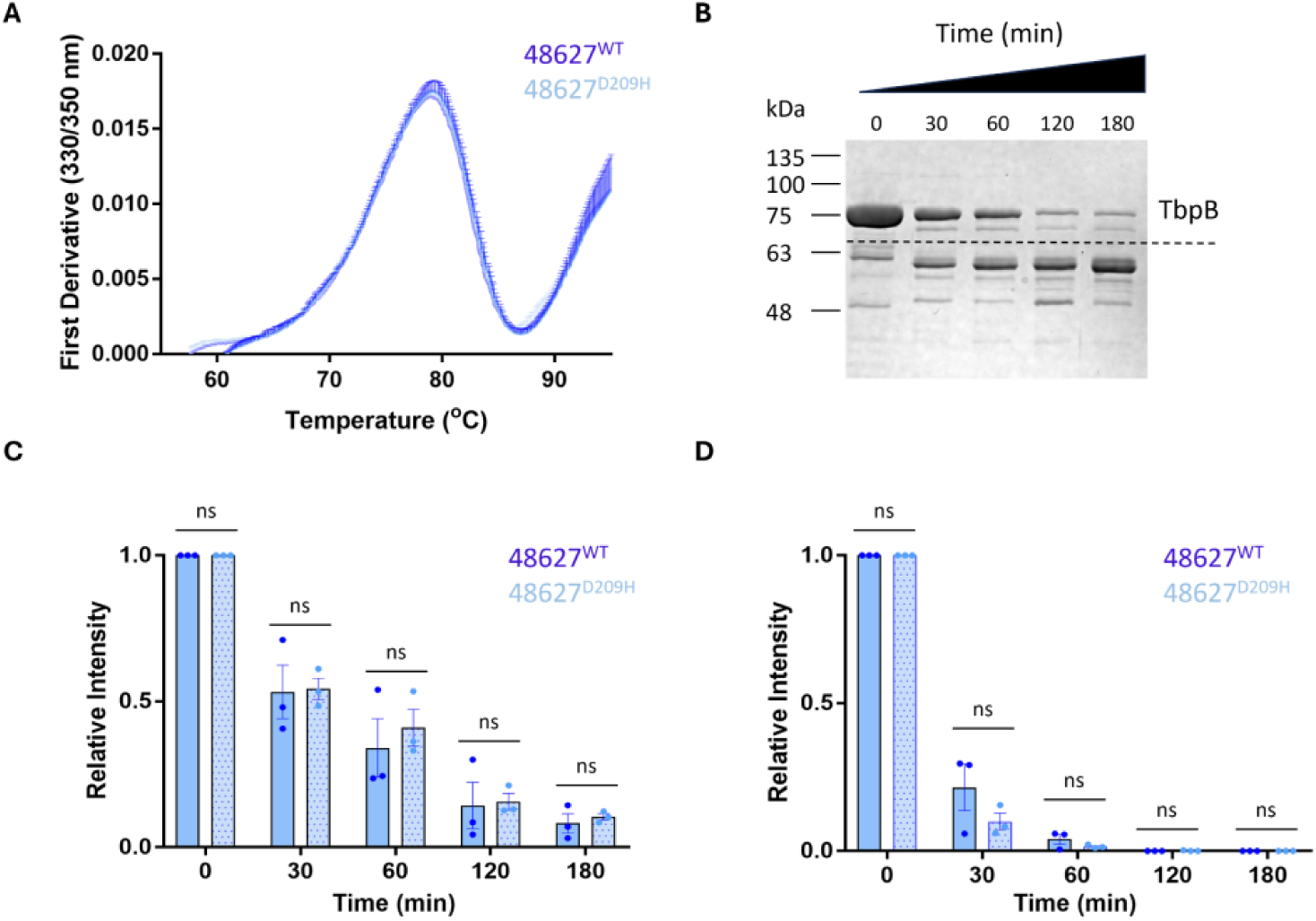
Binding defective mutation does not significantly affect the structural integrity of the 48627 TbpB protein. (**A**) Analysis of thermal stability of the TbpB 48627^WT^ and 48627^D209H^ variants using nano-DSF. (**B**) Trypsin digestion of the 48627^WT^ up to 180 minutes as visualized by SDS-PAGE. Both full-length proteins are approximately 75 kDa, as indicated above the dashed line. (**C**) Quantification of relative intensity of the intact protein over the course of digestion with chymotrypsin and (**D**) trypsin (n=3, *p≤0.05, non-significant (n.s.)). Using ImageJ, the intensity of the full-length protein was normalized to the untreated control (at time 0). Each point represents a single technical replicate and statistically analyzed with an unpaired student t-test.

### Binding-defective mutant TbpB vaccine reduces colonization by *Ngo* in hTf-expressing transgenic mice

To compare the efficacy of each wild-type and mutant antigen pair, we performed lower genital tract challenges in transgenic mice expressing hTf, which is required to consider the contribution of transferrin binding by the pathogenic *Neisseria*^27^. Briefly, female transgenic mice were immunized with either wild-type or mutant TbpB formulated with Alhydrogel or with the adjuvant alone, followed by vaginal infection with ∼10^7^ CFU of the relevant challenge strain of *N. gonorrhoeae*. For our initial study, we immunized and boosted mice with TbpB variants derived from *Ngo* MS11 and then challenged with the homologous strain, since this is the prototype routinely used for mouse studies (Figure 4A). Mice immunized with adjuvant alone had a median colonization of 15 days with the MS11 strain (Figure 4C). The MS11^WT^ antigen reduced median colonization time to 10 days while the MS11^R200E^ mutant antigen further reduced colonization time to a median of 5 days. While the results are consistent with our hypothesis that the mutant would outperform wild type TbpB as an immunogen, the small size of the cohorts prevented them from being statistically significant.

**Figure 4.**
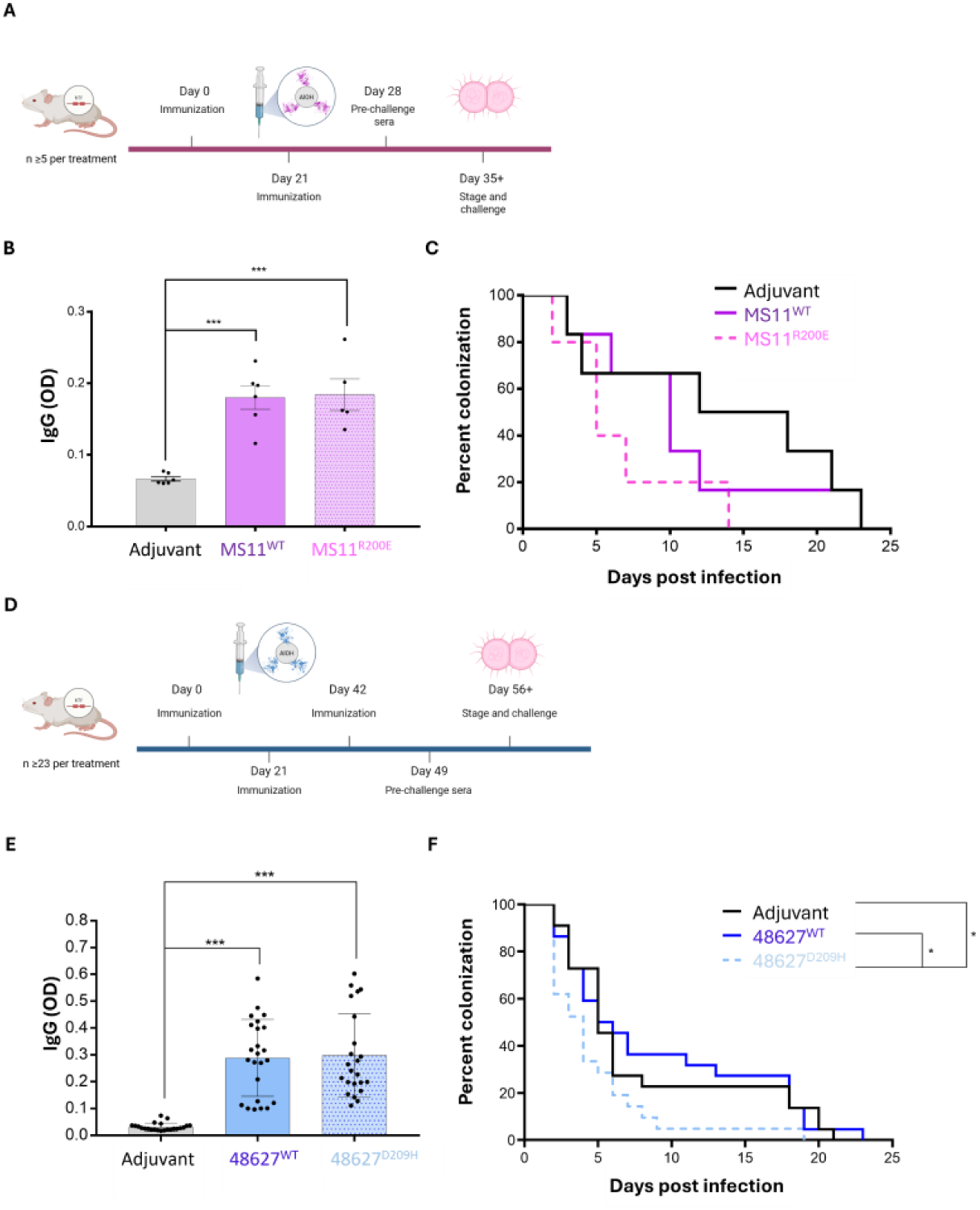
Vaccination with formulations containing mutant TbpB variants leads reduced colonization upon lower genital tract challenge with Ngo. (**A**) Schematic of immunization and challenge with *Ngo* MS11. Female hTf-transgenic BALB/c mice were immunized twice with the Alhydrogel-adjuvanted TbpB prior to challenge. Sera collection prior to challenge is indicated at day 28. (**B**) Quantification of wild-type MS11 TbpB-specific IgG titers at 7 days prior to infection. Each point represents serum from individual mice, analyzed with a one-way ANOVA and *post hoc* Tukey’s test (n≥7, ***p≤0.001). (**C**) Colonization of the immunized mice with *Ngo* MS11 (n≥5). (**D**) Schematic of immunization with TbpB^48627^ and challenge with a matched strain, WHO L. Female transgenic BALB/c hTf mice were vaccinated thrice with the Alhydrogel-adjuvanted TbpB^48627^-derived variants. Prior to challenge (day 49), sera was collected. (**E**) Comparison of the wild-type 48627 TbpB-specific IgG titers of mice immunized with different treatments prior to challenge. Each point represents individual mouse sera, analyzed using a one-way ANOVA with a *post hoc* Tukey’s test (n≥23,***p≤0.001). (**F**) Colonization of the immunized mice with *Ngo* WHO L. Significance in the mouse infection studies was determined through the Mantel-Cox test (n≥21, *p≤0.05).

The 48627 sequence was bioinformatically selected from available genome sequences as a representative of the large phylogenetic cluster of TbpB^23^, but we do not have access to this strain. Thus, we selected a strain from the WHO reference panel that expressed a TbpB sharing a closely related, albeit still heterologous, sequence (WHO L, ∼90% identity)^34^ to test its protective efficacy. For this experiment, we increased the cohort size and immunized mice a third time (Figure 4D). We unexpectedly observed no difference in median colonization of 5 and 5.5 days for immunizing groups of the adjuvant alone and 48627^WT^ immunizations, respectively (Figure 4F). However, immunization with the 48627^D209H^ mutant antigen reduced the median colonization to 4 days and only 1 mouse remained colonized after day 9, which was significantly different from both adjuvant and 48627^WT^ antigen. Thus, for two different gonococcal TbpB variants, a reduction in colonization time was observed when a binding-defective mutant antigen was used as immunogen.

### Antibodies elicited by wild-type and mutant TbpB antigen promote bacterial killing

Given that binding-defective TbpB mutants conferred protection superior to that by their wild-type counterparts, we sought to evaluate the antibody responses generated during our challenge experiments. Prior to the mouse infectious challenge, sera was collected to determine TbpB-specific titers by ELISA with the wild type TbpB protein. No significant differences in average antibody titers between wild-type and mutant TbpB-immunized mice (Figure 4B, E), indicating that the difference in protection was not due to altered immunogenicity of the proteins.

Next, we aimed to measure the functional activity of these antisera. While the iron-regulated expression of TbpB assures that it is expressed *in vivo*, it is not expressed in standard culture medium that is replete with iron. We therefore cultured *Ngo* in low-iron conditions and then verified TbpB expression through immunoblotting before use for these analyses (Supplemental Figure 4). To compare the ability of each antisera to promote complement-dependent killing of gonococci, we performed a serum bactericidal assay. To validate our assay, we also employed the control murine monoclonal antibody 2C7, since this targets a very high density gonococcal epitope on LOS^35^ and effectively activates the complement cascade to promote bacterial killing^36^. We used human serum as a complement source because *Ngo* has the capacity to interact with negative regulators of complement in a host-specific manner^37,38^. We observed a reduction of over 60% in *Ngo* viability after 60 minutes with 2C7 (Figure 5A). Strikingly, antibodies isolated from sera collected from mice immunized with the 48627^D209H^ variant also caused ∼50% reduction in bacterial viability, while those from mice immunized with either 48627^WT^ or adjuvant alone were not bactericidal at the concentrations tested; this unexpected difference in bactericidal activity between the 48627^WT^ and 48627^D209H^-derived antisera was statistically significant.

**Figure 5.**
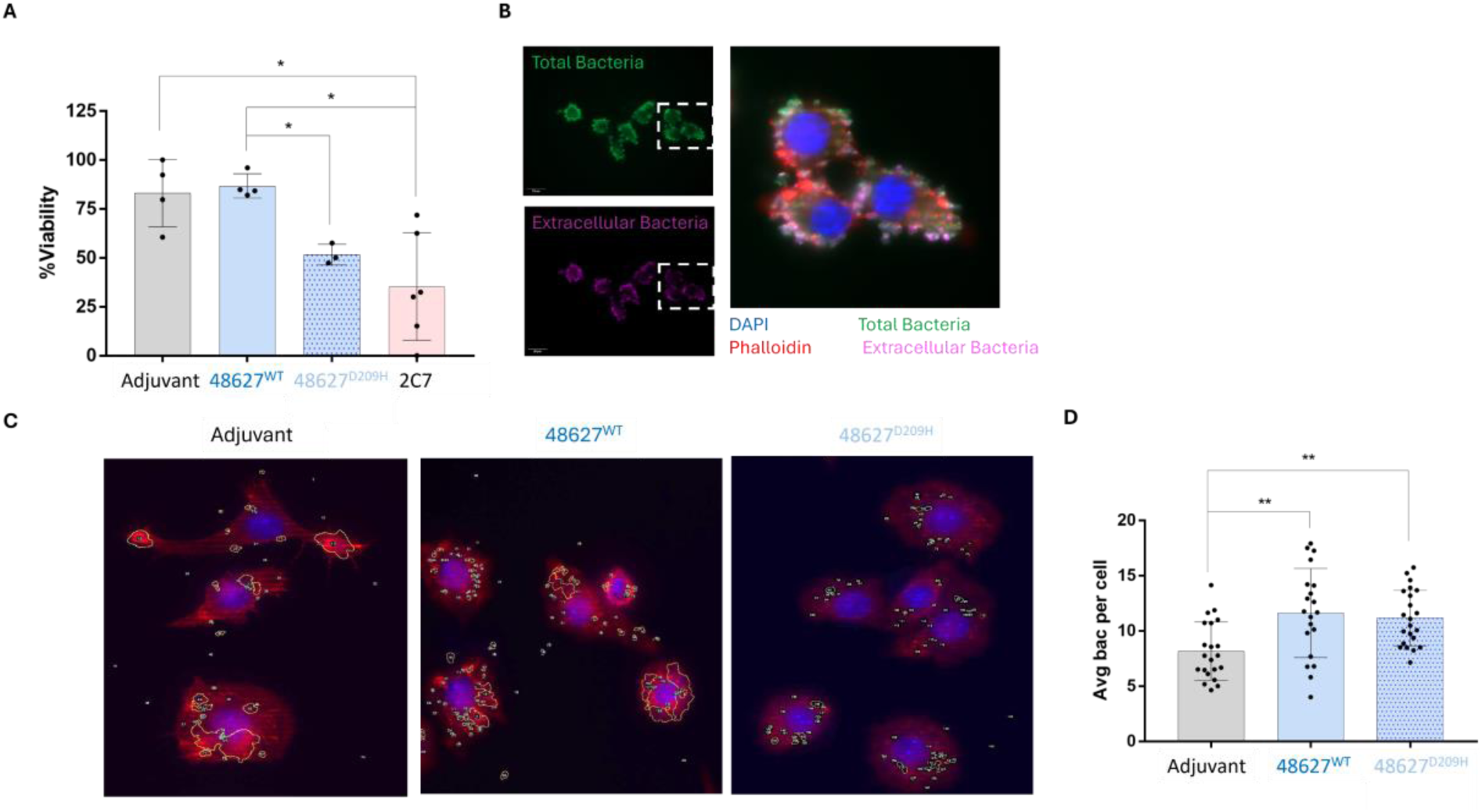
Antisera induced from immunization with 48627^D209H^ antigen promotes bacteria killing. (**A**) Bactericidal activity of antisera pooled from mice immunized with either wild type or D209H mutant TbpB induced killing of the challenge strain, *Ngo* WHO L. Monoclonal antibody 2C7 was used as a positive control, and each point represents a biological replicate. Viability was calculated by comparing the number of colonies quantified at 0 and 60 minutes following the addition of human complement, with significance analyzed by a one-way ANOVA with a *post hoc* Tukey’s test (n≥3, *p≤0.05). **(B)** Representative image showing infection of RAW macrophages with opsonized WHO L, as visualized by fluorescence microscopy. Gonococci were opsonized by antisera derived by pooling sera from mice immunized with adjuvant, 48627^WT^ or 48627^D209H^ for 30 minutes and then incubated with macrophages for 60 minutes. Macrophages were stained to distinguish nuclei and phalloidin, while bacteria were stained before and after permeabilization to differentiate between total (green) and extracellular (purple) bacteria. Scale bar represents 50 µm. **(D)** Visualization of internalized gonococci within murine macrophages by fluorescence microscopy. At least 5 images obtained from each of 3 or more coverslips produced from at least two biological replicates. Internalized bacteria are emphasized by white arrows. Images below illustrate regions quantified as total bacteria as identified by Fiji. (**E**) Quantification of opsonized gonococci associated with RAW macrophages was performed using Fiji (Image J) (Supplemental Figure 5) (n_images_≥19, n_macrophages_≥200, **p≤0.01).

To compare the capacity for these antisera to promote phagocytic clearance of the bacteria, we performed opsonophagocytic assays with RAW 264.7 cells, a murine-derived macrophage cell line^39^. The *Ngo* WHO L strain was exposed to antisera from mice that had been immunized with either 48627^WT^ or 48627^D209H^ TbpB variants, and these were then applied to the macrophage cultures. We observed that antisera from mice that had been immunized with the mutant TbpB variant increased bacterial binding and engulfment using fluorescence microscopy after differential staining of intracellular and extracellular *Ngo*. While there appeared to be a more reproducibly higher number of *Ngo* per cell with the mutant TbpB-derived antisera, this was not significantly different than the sera elicited by the wild type TbpB vaccine (Figure 5B,C,D). When considered together, both TbpB variants elicit a robust humoral response that can promote phagocytic clearance of *Ngo*, but complement-dependent killing was significantly more effective when the bacteria were exposed to antisera from animals immunized with the 48627^D209H^ mutant variant.

### Binding-defective mutant immunogen elicits a greater titre of TbpB-neutralizing antibodies

In considering the underlying mechanism to explain the enhanced effectiveness of our vaccine based upon the mutant TbpB at reducing gonococcal colonization, we postulated that the mutant variant would avoid complexing with the hTf present within tissues, thereby allowing it to elicit more antibodies targeting the hTf-binding interface of TbpB. If true, then the antisera from 48627^D209H^-immunized animals should be more effective at blocking hTf binding than wild type TbpB. Therefore, using a protein-based assay, we assessed hTf binding to wild type TbpB in the presence of antisera from the three different immunized groups. We consistently observed that the mutant variant elicited higher hTf blocking titers in hTf-expressing transgenic mice relative to the wild-type antigen and adjuvant control for both variants (Figure 6A,B). In contrast, the wild-type antigen could only consistently raise function-blocking antibodies in wild type mice, but not transgenic, inferring that hTf binding affected the antibody response (Supplemental Figure 5).

**Figure 6.**
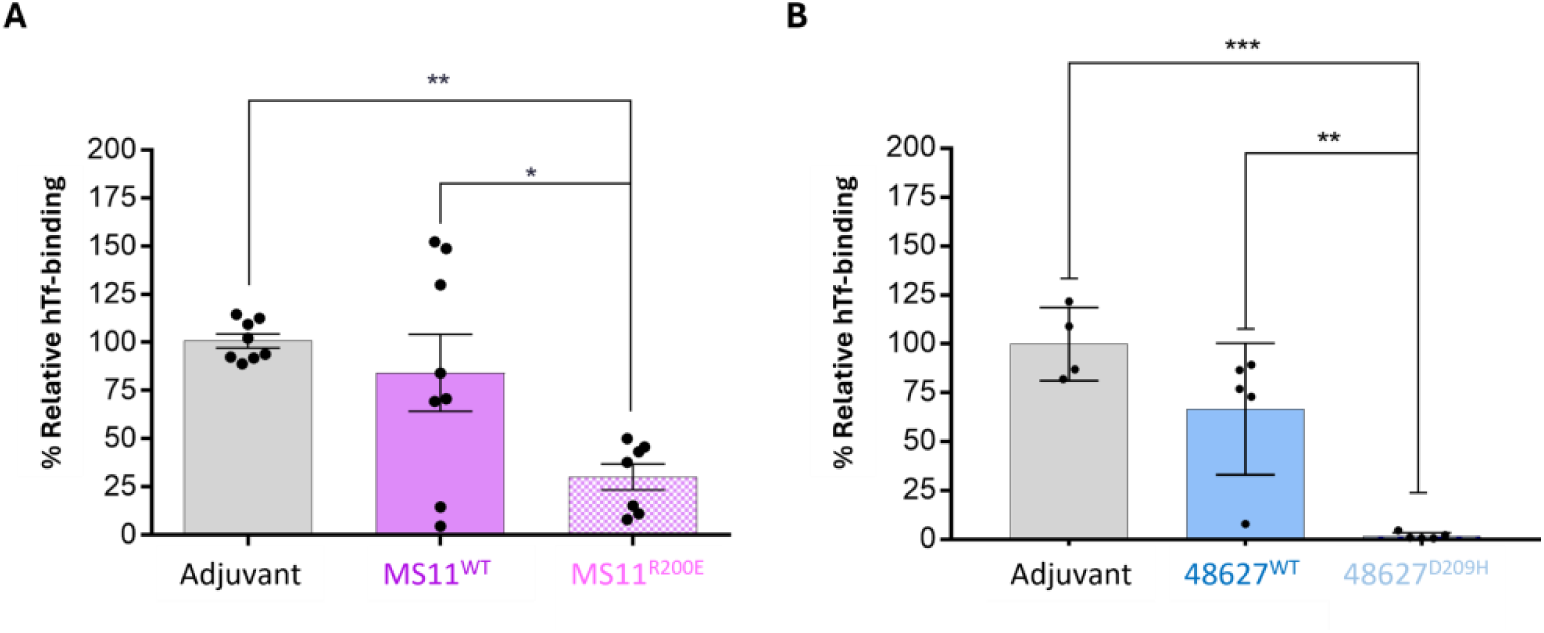
Antibodies elicited by binding-defective mutants more effectively block hTf binding. (**A**) Relative amount of hTf binding determined through protein-based binding assay using purified TbpB in the presence of different sera. Transgenic BALB/c hTf mice were immunized with either adjuvant, MS11^WT^, or MS11^R200E^ TbpB. Binding was calculated in reference to a no antibody control included in each technical replicate. Each data point represents individual mouse sera diluted 1:50, and statistically analyzed using a one-way ANOVA with a *post hoc* Tukey’s test (n≥7, *p≤0.05,**p≤0.01). (**B**) hTf blocking in the presence of sera from transgenic mice immunized with adjuvant, 48627^WT^, or 48627^D209H^ mutant (n≥3, **p≤0.01, **p≤0.001).

## 3. Discussion

While subunit vaccines are often considered to be inert when expressed in isolation, components that have a simple binding function retained within the vaccine preparation may engage with their host target. While this would normally be considered by experimental studies in animals where the binding interaction is conserved, it can not occur when the binding function is restricted to targets unique to the pathogen’s natural host. This is a particular challenge with pathogens such as *Ngo* and *Nme*, which are highly evolved to life within humans, since virulence factors frequently only bind the human form of their targets^6^. The potential implications of this have been shown previously with vaccines based upon the meningococcal FHbp^26^ and NspA^6^, both of which bind specifically to the human complement component, factor H. While these proteins are both effective at eliciting a protective response in wild-type mice, their efficacy as a vaccine is reduced in transgenic mice that express human factor H. We have observed a similar effect when immunizing pigs with TbpB from the pig-restricted pathogen, *Glaesserella parasuis* (formerly *Haemophilus parasuis*)^25^. In this case, a binding-defective mutant of TbpB elicited more robust T-cell and B-cell responses, and superior protection than did the wild-type variant. In this current study, we use complementary protein engineering and mouse genetic approaches to extend this work to human transferrin (hTf)-specific TbpB proteins from *Ngo*, establishing that point mutations that abrogate binding to hTf confer superior protection against lower genital tract infection by this sexually-transmitted bacteria.

The bacterial transferrin receptor is considered an obvious vaccine target for a variety of different host-restricted bacterial pathogens, including the gonococcus^22–25,40–47^. The lipid-anchored TbpB is highly accessible at the bacterial surface, a requisite of its transferrin-binding function, and is a stable and highly immunogenic protein^23–25,40,48,49^. We have established that wild-type mice immunized with gonococcal TbpB develop a broadly cross-reactive humoral response and are protected against infection by *Ngo*^23,24^. We have taken advantage of transgenic mice expressing hTf to determine whether TbpB-based vaccine efficacy could be influenced by hTf binding. We have used a combination of *in silico* modelling based upon the solved structure of other (non-gonococcal) TbpBs and our crystal structure of a gonococcal TbpB N-lobe to guide the rational design of binding-defective but structurally- and immunologically-intact TbpB variants. With these, we found that a vaccine comprised of the binding-defective TbpB elicited similar titres of antibody but was significantly more effective than the wild-type variant at protecting hTf-expressing mice. This effect may result from the propensity for native TbpB binding to hTf once the vaccine is administered to the tissues, which would sterically prevent this functional surface from engaging with B cells to elicit an adaptive humoral response. Unexpectedly, we also observed that the binding-defective mutant elicited a more bactericidal response. It is enticing to consider that this may result from increased epitope density when binding and non-binding surfaces of TbpB are exposed, however the formal testing of this mechanistic premise remains beyond the scope of this study.

While the TbpB structure is available for several other species, these consistently indicate that most sequence and structural variation occurs within their surface loops^50^. This suggests that the functional loops undergo selective pressure to evade immune responses mounted by the host, contributing to the antigenic diversity of TbpBs. Consistent with this, AlphaFold3 predicted the canonical folding of *Ngo* TbpBs similar to other surface lipoproteins, but all loops containing functional (hTf-binding) residues on the N-lobe were predicted with lower confidence^29^. Despite this sequence diversity, there appears to be a pressure to maintain charged amino acids at similar spatial positions. As seen in meningococcal TbpBs from each isotype, represented by M982 and B16B6 TbpB variants, a positively charged residue was maintained (R211 and K194, respectively,from PDB 3VE2 and 4QQ1)^20,51^. Similarly, mutation of the gonococcal MS11 TbpB R200 residue was found to significantly influence binding to hTf. Beyond *neisserial* TbpBs, the TbpB variants from the porcine pathogen *A. pleuropneumoniae* also have a positively charged residue, R179, implicated in transferrin binding (PDB: 3PQS)^52^. We have exploited this conservation across a diverse group of TbpBs to identifying potential functional residues on computational models generated for uncharacterized TbpB variants, accelerating mutant design and vaccine development. In the context of *Ngo*, we modelled and engineered mutations that disrupt hTf binding by the MS11 and 48627 TbpB variants, and found both mutants to confer more effective protection against gonococcal colonization compared to the wild type proteins when using hTf-expressing transgenic mice.

While our work found that the binding-defective TbpB mutants produced superior vaccines, previous studies indicate that mutant antigens do not universally improve efficacy. Previously, two different binding-defective TbpB mutants were tested in a *G. parasuis* tracheal challenge model in pigs^53^. Only one of these increased survival of the vaccinated pigs (80%) after the infectious challenge^53^. Both antigens elicited antibodies that bound intact bacteria, initiated complement deposition, and stimulated cellular immune mechanisms^53^. This suggests that there are characteristics beyond the inability to bind transferrin that contribute to the increased protective efficacy observed in an earlier heterologous challenge^25^. While the study evaluated the immunogenic potential of both mutants in *G. parasuis,* it lacked an in-depth analysis of the structural consequences of the point mutation and its influence on immune activation^53^. For this reason, we thoroughly investigated the biophysical and immunological characteristics of mutants relative to their respective wild-type variants.

*In silico* modelling for both mutated gonococcal TbpB antigens predicted that the point mutations would not affect the overall folding of the protein, maintaining other key conformational epitopes found on TbpB. Beyond modelling, we compared the thermal stability and resistance to protease degradation of the recombinant wild type and mutant derivatives, provide insight into the structural integrity of the proteins^54^. For the 48627 variant, nano-DSF verified that the mutation did not affect the thermal stability between the wild-type and mutant proteins, and limited proteolysis with enzymes chymotrypsin and trypsin, serine proteases that target different peptide sequences, had comparable effects on the 48627^WT^ and 48627^D209H^ proteins (Figure 3B). The MS11 variant mutant, MS11^R200E^, also had no differences in the secondary structure or size when analyzed by Far-UV CD analysis, suggesting that it was also structurally intact. While it must be recognized that the biophysical characteristics and enzyme susceptibility may change upon protein adsorption to the adjuvant^55^, our results demonstrate that the point mutations that we introduced did not impact the structural integrity of TbpB, despite abrogating hTf binding.

In transgenic mice expressing hTf, we showed that mutant TbpB elicits greater protection compared to wild type TbpB. For both TbpB mutants, we observed an increased clearance of *Ngo* in mice immunized with binding-defective TbpB antigens compared to wild type protein, with a significant difference observed between the 48627^WT^ and the 48627^D209H^ antigen. While the wild type MS11-derived TbpB appeared to accelerate clearance of the *Ngo* relative to adjuvant alone, this was less effective than the mutant protein. In contrast, mice immunized with the wild-type 48627-derived TbpB had no benefit over the adjuvant alone, while the mutant variant of this protein allowed all but 1 mouse to clear the infection. It is important to note that this variant is effective in the absence of hTf since we previously reported that immunization with 48627^WT^ TbpB protected wild type C57BL/6 mice against the same WHO L strain^23^. These results implicate binding to hTf in diminishing the effectiveness of the immune response, and underscores the importance of employing appropriately selected animal models to support the development of effective prophylactics targeting human pathogens like the gonococcus.

Despite the variability within its hTf binding loops, we previously identified variants capable of inducing a cross-reactive antibody response. The 48627 variant was selected to represent the largest phylogenetic cluster of TbpBs, and we subsequently validated its ability to elicit a broadly cross-reactive antibody response, allowing development of a bivalent TbpB vaccine that confers broad coverage against *Ngo*^23^. Our current work highlights that rationally-designed binding-defective mutants of these TbpB components must be used to avoid hTf complexing of the vaccine components when administered to humans. In a broader sense, this reinforces the importance of accounting for host-pathogen interactions in vaccine design^56,57^.

## 4. Supplemental Figures

**Supplemental Figure 1.**
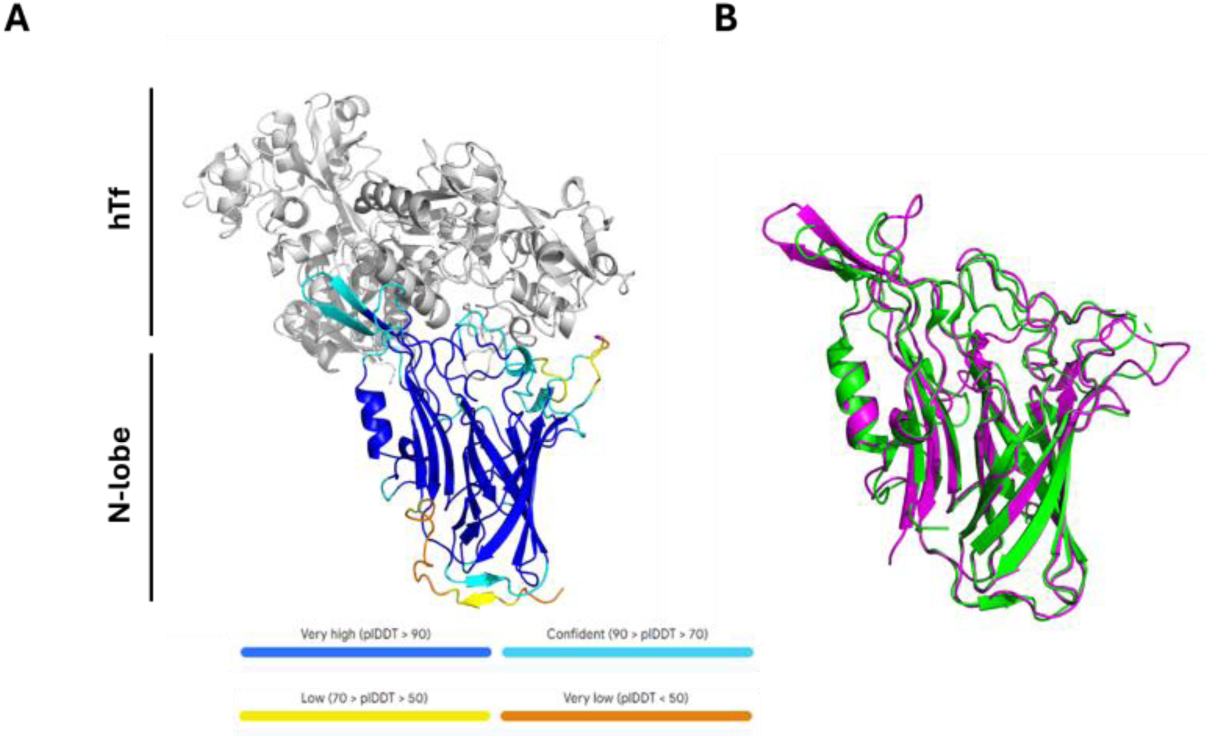
Structural alignment of Ngo MS11^WT-Nlobe^ to M982 TbpB and AlphaFold predicted models. **(A**) AlphaFold3 model showing hTf (grey) complexed to MS11^WT-Nlobe^. The structure of hTf was predicted with high confidence, with a pIDTT>80 and is shown in grey. Similarly, the β-barrel and handle domains, which is canonical in other TbpBs, was also predicted with high confidence. In contrast, orange represents regions predicted with lower confidence. **(B)** Overlay of MS11^WT-Nlobe^ to M982 TbpB N-lobe. The meningococcal M982 TbpB (PDB:3VE2) N-lobe is shown in green and MS11^WT-Nlobe^ is shown in violet^20^. Alignment shows similar backbone structure with more variability in the loop region.

**Supplemental Figure 2.**
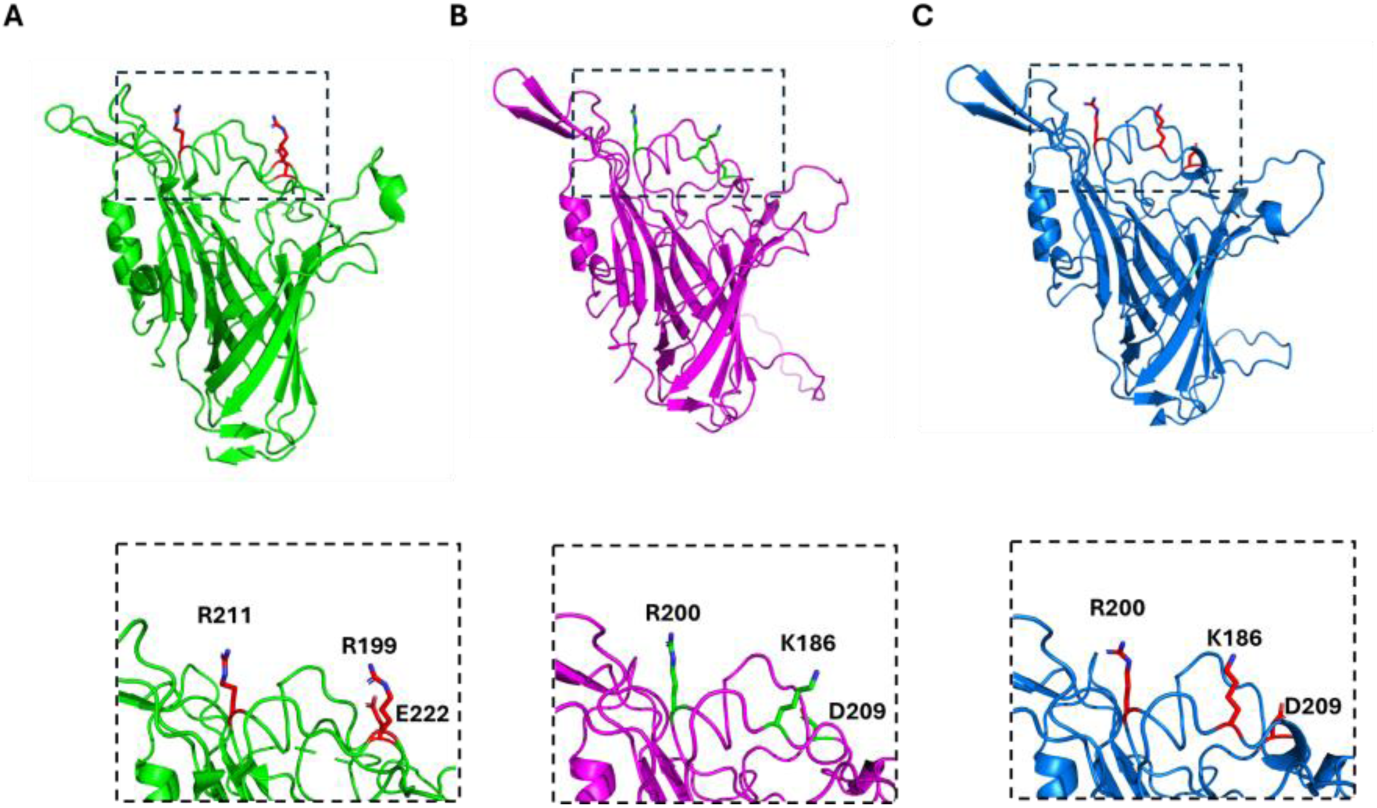
Residues on TbpB N-lobe involved in hTf binding. **(A)** Crystal structure of meningococcal M982 TbpB (PDB: 3VE2). Residues implicated in hTf binding, as inferred from structure and mutagenesis studies, are indicated in red. **(B)** AlphaFold3 prediction of the gonococcal MS11^WT^ and **(C)** 48627^WT^ TbpB N-lobes. Through structural alignment to the M982 TbpB, three residues on each TbpB that may be potentially important for function are represented in green (MS11^WT^) and red (48627^WT^).

**Supplemental Figure 3.**
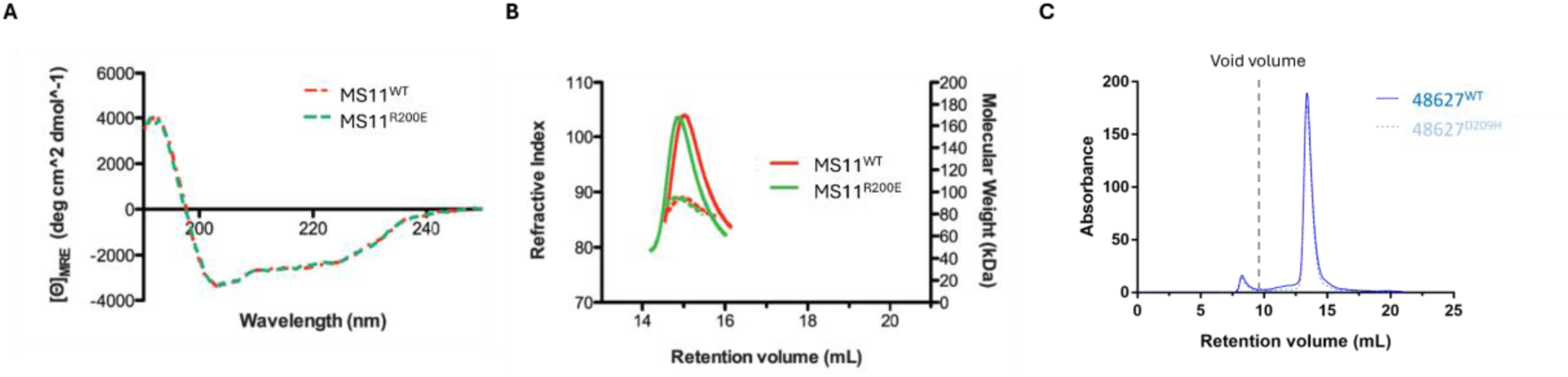
Biophysical characterization of selected TbpBs and their respective mutants. **(A)** Far-UV CD analysis of MS11^WT^ and MS11^R200E^ between 190 to 250 nm. **(B)** Size exclusion chromatography-multi-angle light scattering of MS11^WT^ and MS11^R200E^. **(C)** Size exclusion chromatography of the 48627 antigens. Approximately 1.25 mg of each protein was injected onto a Superdex 200 Increase 10/300 GL in PBS.

**Supplemental Figure 4.**
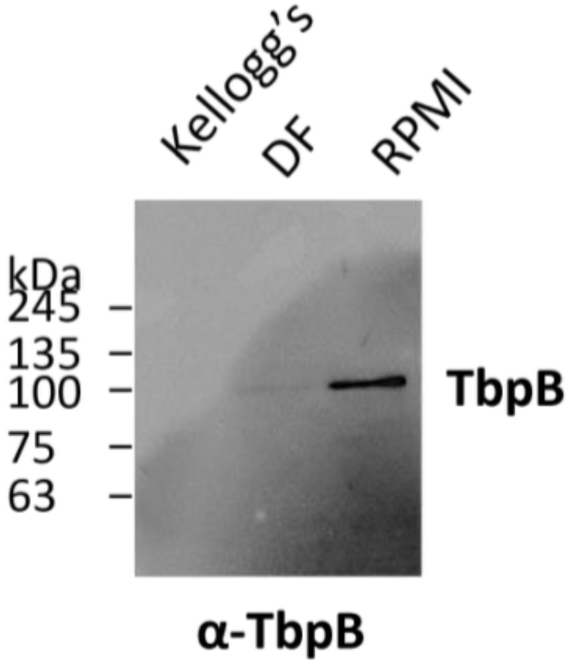
Comparison of TbpB expression upon growth in different culture media. To optimize TbpB expression, the WHO L strain was grown overnight on solid GC Kellogg’s media at 37°C with 5% CO_2_. Using a Dracon swab, bacteria were replated onto a GC Kellogg’s plate supplemented with 10 µM deferoxamine mesylate (DF) and grown for 4 hours under the same conditions. In contrast, bacteria were resuspended in iron-restricted RPMI 1640 overnight (RPMI) at 37°C with agitation at 160 rpm for 16 hours. A TbpB-specific antisera from immunizations with 48627^WT^ was used to probe for TbpB expression.

**Supplemental Figure 5.**
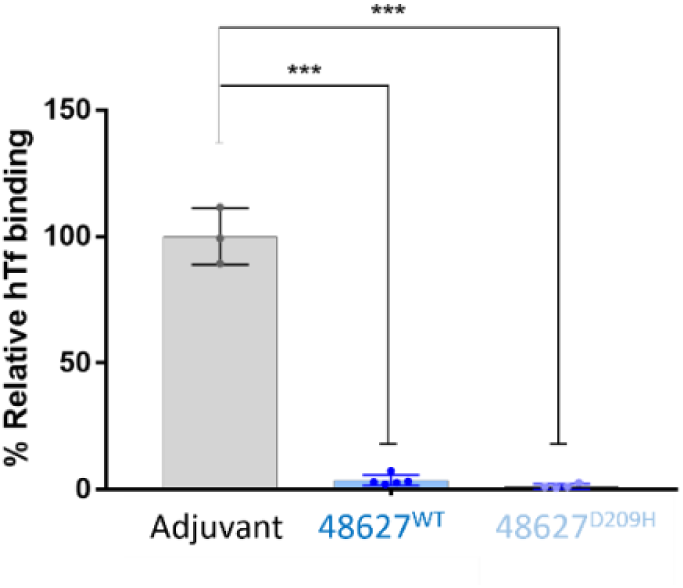
Antibodies elicited from immunization in wild type mice with either wild-type or mutant TbpB consistently block hTf binding. Wild-type BALB/c mice expressing mouse transferrin were immunized thrice with either adjuvant, 48627^WT^, or 48627^D209H^. In a protein-based immunoassay, sera was incubated with homologous TbpB before the addition of hTf. Each point represents individual mouse sera, with statistical analysis by one-way ANOVA (n≥3, ***p≤0.001).

## 5. Methods and Materials

### Antigen expression and purification

MS11^WT^ and MS11^R200E^ *tbpB* genes were subcloned into a pET28a vector in fusion with an N-terminal histidine tag for cytoplasmic expression. The R200E mutant and two SERp mutants were generated by quick change mutagenesis (see Supplementary Methods). Full length protein for each construct was expressed and purified from *Escherichia coli* ER2566 (New England Biolabs, cat. C2566), cells were grown in an overnight starter culture in LB at 37°C with shaking at 175 rpm. The starter was subcultured 1:100 into fresh LB media supplemented with kanamycin (50 µg/mL) and grown for 4 hours at 37°C with shaking (175 rpm) and then protein expression was induced by the addition of 250 µM Isopropyl β-D-1-thiogalactopyranoside (IPTG) for a further 16 hours shaking (175 rpm) at 20°C. Cells were harvested (5000x*g*, 30 minutes) and resuspended in lysis buffer (50 mM Tris pH 8, 300 mM NaCl, 10 mM imidazole) and lysed by sonication (Branson S450). The lysate was clarified by centrifugation (16000x*g*, 90 minutes) and syringe filtering (0.2 µm) before loading on NiNTA resin (ThermoFisher Scientific). His-tagged TbpB protein was eluted from the resin, buffer exchanged into a low salt buffer and then injected over a MonoQ 5/50 GL anion exchange column (Cytiva) through an ÄKTA purifier unit. The sample was washed and eluted by a salt gradient ranging from 50 mM to 2000 mM NaCl buffered with 50 mM of Tris at pH 8. Protein was collected and concentrated prior to being flash frozen and stored at -80°C until crystallization or immunization.

The 48627 TbpBs were secreted into media using its cognate translocon, the surface lipoprotein assembly modulator (SLAM)^58^, thereby making antigen preparation more efficient by reducing the number of purification steps. The 48627^WT^ and 48627^D209H^ *tbpBs* were subcloned into a vector that co-expresses the neisserial SLAM protein^58^ by restriction-free cloning. With SLAM constitutively expressed, the his-tagged TbpB, which was under the control of a rhamnose promotor, was secreted into media upon the addition of the inducer in a SLAM-dependent manner. Briefly, *E. coli* C43 was transformed with the secretion constructs and plated onto LB agar with kanamycin selection for growth overnight at 37°C with shaking at 175 rpm. Multiple colonies were grown in small volumes of LB containing the kanamycin for up to 6 hours at 37°C with shaking and subsequently sub-cultured into a larger volume for growth in same conditions for 12 hours. 0.4% rhamnose (BioShop, cat. RHA001) was added for TbpB expression and the culture was grown for an additional 3 hours at 37°C with shaking (175 rpm) and then harvested (5000x*g*, 30 minutes). The resulting supernatant was filtered (0.22 µm) and circulated through a HisTrap EXCEL (Cytiva, cat. 17371205) column overnight at 4°C to capture the protein of interest. The column is subsequently washed with buffer (50 mM Tris pH 8, 300 mM NaCl) and eluted with imidazole (50 mM Tris pH 8, 300 mM NaCl, 300 mM imidazole pH 7.4). The eluted fractions were collected and dialyzed into phosphate buffered saline (PBS) for 1 hour at 4°C. Thrombin (Sigma-aldrich, cat. T8885) was added to the dialyzed protein to cleave the his-tag and further dialyzed in PBS overnight at 4°C. Next, p-Aminobenzamidine beads (Sigma-aldrich, cat. A7155) were added and incubated for 3 hours to bind and remove the thrombin with gentle rocking at 4°C. Beads were removed through low-speed centrifugation and syringe filtration (0.2 µm) and protein was collected and concentrated prior to being flash frozen and stored at -80°C until use.

### Protein crystallization, data collection and structure determination

Purified MS11 and 48627 TbpB did not crystallize in over a thousand screening conditions. Due to challenges in crystallizing the full-length protein, the Surface Entropy Reduction prediction (SERp) server was used to identify clusters of high entropy residues for potential targeted mutagenesis to aid in crystallization^28^. Two high-entropy conserved clusters (K218, K219, E220) and (E236, K237, K238) were identified for both MS11^WT^ and 48627^WT^ TbpBs and referred to as SER1 and SER2. Since functional residues were in the N-lobe of TbpB, we generated an N-lobe only construct, consisting of residues 19 to 361 of the mature MS11^WT^ protein with the two SERp clusters incorporated to generate MS11^WT-Nlobe^. Both clusters were mutated into alanine residues to enhance crystallization on the MS11^WT-Nlobe^. A similar workflow to the purification of the full-length protein was used to produce the MS11^WT-Nlobe^. For structural characterization of MS11^WTN-lobe^, the protein was dialyzed to a lower salt concentration (50 mM NaCl) following affinity chromatography and during anion-exchange, the Mono Q resin was washed with a buffer with lower salt (300 mM NaCl). Protein was collected and concentrated prior to being flash frozen and stored at -80°C until immunization.

Purified MS11^WT-Nlobe^ at 9 to 10 mg/mL was screened against multiple crystallization screens (Hampton Research, Qiagen, Microlytic, Anatrace) by sitting drop vapour diffusion in 96-well Intelli-Plates using a Gryphon drop setter (Arts Robbins Instruments) using 1:1 protein:precipitant ratio. Optimization screens in 24-well sitting drop Intelli-Plates were set up with drops in 2:1 and 1:1 protein:precipitation ratio at room temperature. Microseeds from crystals obtained in the initial optimization screens were prepared using seed beads (Hampton Research). Diffraction quality crystals for MS11^WT-Nlobe^ was successfully optimized and obtained in 0.1 M sodium calcodylate pH 6.5 and 1 M sodium citrate and flash frozen in liquid nitrogen for structural determination at the synchrotron.

X-ray diffraction data were were collected on the 08ID-1 beamline at the Canadian Light Source (CLS, Saskatoon, Saskatchewan, Canada) at a wavelength of 0.9795 Å with 0.2° oscillations, totaling 900 images^59^. The diffraction data was integrated and scaled using XDS to a resolution of 2.59 Å^60^. The crystal structure was solved through Phenix (ver. 1.21.2-5419) using Phaser molecular replacement with an AlphaFold 2 predicted model of the MS11^WT^ TbpB^61,62^. It was refined using Coot (ver. 0.9.8) and Phenix Refine at a resolution of 2.59 Å^63,64^. The diffraction data and refined coordinates are deposited in the Protein Data Bank with PDB code 9Q07. PyMol (Schrödinger LLC) was used to visualize and generate structural figures^65^.

### Solid Phase Binding Assay

For small-scale expression of protein, each TbpB was cloned into a custom vector encoding an N-terminus his tag, followed by a biotin acceptor peptide (BAP), and a maltose binding protein (MBP) tag to aid purification and solubility. Similar to the cytoplasmic antigen expression described above, the plasmid was transformed into *E. coli* ER2566. Following recovery in LB, transformed bacteria were diluted 1:1000 and grown in ZY-5052 auto-induction media ((10 g tryptone, 5 g yeast extract), 50 mL NPS (3.3 g ammonium sulfate, 6.8 g potassium phosphate monobasic, 7.1 g sodium phosphate dibasic), 20 mL 5052 (5 mL glycerol, 0.5 g glucose, 2 g lactose) and 1 mL magnesium sulfate per liter) supplemented with ampicillin (50 µg/mL) at 37°C with 175 rpm overnight^66^. Bacteria were harvested through centrifugation and pellets were resuspended in 50 mM Tris, pH 8.0 and 300 mM NaCl, supplemented with lysozyme (10 mg per L of culture), DNase I (0.8 mg per L of culture), and phenylmethylsulfonyl fluoride (BioBasic, cat. PB0425). Glass beads were added and used to lyse cells via vortexing, with lysate clarified through high speed centrifugation (21300x*g*, 10 minutes). Lysate was incubated with Ni-NTA resin and eluted using imidazole-containing buffer (50 mM Tris, pH 8.0, 150 mM NaCl, 300 mM imidazole, pH 7.4). Protein was dialyzed in phosphate buffered saline (PBS) prior to application in subsequent assays.

Under 5 µg of the purified protein (1 mg/mL) was applied to a nitrocellulose membrane and subsequently blocked with 5% skim milk in PBS for 1 hour at RT. The blot was washed with PBS with 0.1% Tween 20 (BioShop, cat. TWN508) (PBS-T) thrice. To infer binding, membranes were incubated with hTf conjugated to HRP (Rockland, cat. 009-0034) for 1 hour at room temperature (RT) and washed with PBS-T thrice. Lastly, membranes were incubated with Enhanced Chemiluminescence substrate (BioRad, cat. 1705061) at RT in the dark and visualized.

### Biolayer interferometry

Kinetic experiments were performed using the Octet RED96 FortéBio system at RT in a similar manner as reported previously^67^. Streptavidin sensors (Satorius, cat. 18-5019) were activated in PBS with 0.1% bovine serum albumin (BSA, BioShop Canada, cat. ALB001) (kinetics buffer) and loaded with purified his-tagged-TbpB for 60 seconds. Loaded probes were dipped into increasing concentrations of holo-hTf (Sigma, cat. T0665-1G) suspended in PBS for 200 seconds and dissociated in kinetics buffer for the same time. Wild-type protein was tested up to 300 nM hTf and mutants, which did not demonstrate binding at these concentrations, were tested up to 10000 nM hTf. Using the association and dissociation curves generated, *K_D_* was determined with kinetic and equilibrium analysis using GraphPad Prism (version 7.03).

### Nano-DSF

Using NanoTemper Tycho NT.6, the proteins were loaded into capillaries (NanoTemper, cat. TY-C001) at a concentration of 0.1 mg/mL in PBS^68^. The absorbance of the proteins were measured at 330 and 350 nm at a range of temperatures, up to 95°C for three technical replicates. The inflection temperature, which infers melting temperature, was estimated based on the ratio of absorbance at 330 and 350 nm.

### Limited Proteolysis Assay

TbpB proteins were diluted in PBS to a concentration of 0.2 mg/mL for a total volume of 60 µL. Chymotrypsin (0.6 µg, Promega, cat. V1061) or trypsin (0.3 µg, Promega, cat. V5111) was added to protein and incubated at 37°C. 10 µL of each reaction was added to an equal volume of 2X SDS buffer containing β-mercaptoethanol (BME) were taken at 30 minutes until 180 minutes. Samples were heated at 95°C for 5 minutes and 8 µL was loaded into each well of a 12% SDS-PAGE gel. The gel was stained with Coomassie blue and destained prior to visualization. Densitometry was performed using Image J, normalized to the intensity of the full-length untagged protein, approximated at 75 kDa, at time 0^69^.

### Mice

Mouse studies were performed under the animal use protocol 20011775, approved by the Animal Care Committee at the University of Toronto, which must confer to the ethical and legal requirements of the province of Ontario’s Animals for Research Act and the federal Canadian Council on Animal Care (CCAC). Female homozygous transgenic mice BALB/c expressing hTf, which contains the transgene under a ubiquitous promotor, were bred in-house for immunization and challenge studies^27^. Female wild type BALB/c mice, expressing mouse transferrin, were purchased from Charles River and allowed to acclimate for 1 week before experiments. Only mice over the age of 6 weeks were used in the studies described. All mice were housed in specific pathogen free conditions and were provided food and water *ad libitum.* ^27^

### Immunization

Transgenic female BALB/c mice expressing hTf mice were randomized into receiving either the wild-type or mutant antigen vaccine through intraperitoneal administration. The antigen was formulated in sterile PBS, adjuvanted with aluminium hydroxide (2% Alhydrogel adjuvant, InvivoGen, cat. 21645-51-2) in a total volume of 100 µL. Mice were immunized twice with 20 µg of the MS11^WT^ or MS11^R200E^ mutant, or the adjuvant alone, on day 0 and day 21 in preparation for the (homologous) challenge with *N. gonorrhoeae* MS11. Saphenous bleeds were performed on day -1 (naïve), 20 (primary), 28 (boost, pre-challenge), and cardiac puncture was performed upon euthanasia to collect sera. In contrast, mice were immunized thrice with 15 µg of the 48627^WT^ or 48627^D209H^ mutant or adjuvant control at three-week intervals in prior to infection with *N. gonorrhoeae* WHO L in a (heterologous) challenge. Similarly, serum was collected 2-3 days prior to immunizations and one week before challenge, where they were stored at -20°C for analysis in assays. All immunizations were well-tolerated, with no adverse reactions were observed.

### Lower genital tract gonococcal challenge

For the lower genital tract challenge, female transgenic hTf mice were challenged two weeks following either the second dose to the homologous challenge or the third dose of vaccine for the heterologous challenge. As described previously, mouse vaginas were gently washed with PBS with calcium and magnesium (PBS+/+, Wisent, cat. 311-011) and lavages were examined under light microscopy to determine the stage of the estrus cycle^70^. Based on cell morphology, mice that were in diestrus, denoted by a dense population of neutrophils, were given β-estradiol (0.5 mg in 200 µL sub-cutaneous, Sigma-Aldrich, cat. E4389) to halt them in the estrus phase two days before infection. In addition, antibiotics were administered intraperitoneally (0.6 mg vancomycin (BioShop Canada, cat. VAN990.5) and 2.4 mg streptomycin (BioShop Canada, cat. STP101.5) and were halted following infection. Water with trimethoprim (0.04 g/100 mL, Sigma-Aldrich, cat. T7883) was given to limit growth of commensals.

Two days after b-estradiol administration (day 0) mice were infected intra-vaginally with approximately 10^7^ CFU of *Ngo* WHO L (Public Health England) in which streptomycin resistance was induced for compatibility with this model or mouse-passaged *Ngo* MS11. Bacteria were grown overnight on GC agar (BD, cat. BD228950) supplemented with in-house Kellogg’s or Isovitalex (BD, cat. B11876) at 37°C with 5% CO_2_. Bacteria were collected with a Dracon swab and resuspended to the appropriate concentration, as estimated by OD_550_, in PBS+/+.

Additional doses of β-estradiol were administered subcutaneously on the day of infection and on day 2 post-infection to lock mice in estrus. Antibiotics were injected into mice before and after infection (day -1 to day 1). Mice were administered 400 µL of PBS+/+ subcutaneously following infection to prevent and treat dehydration. Moreover, trimethoprim (0.04 g/100 mL) with streptomycin (0.5 g/100 mL) was added to drinking water to 2 days (day 2) post-infection. Infected mice were lavaged daily (15 µL PBS+/+, diluted into 60 µL) and lavages were diluted up to 1:1000 onto GC agar supplemented with Kellogg’s and VCNT (vancomycin, colistin, nystatin, and trimethoprim, BD BBL^TM^ VCNT Inhibitor (Thayer and Martin), Fisher Scientific, cat. B12408). Mice that did not have any recoverable *Ngo* for three consecutive days were considered to have cleared the infection. Once mice had cleared the infection, they were humanely euthanized by CO_2_ overdose followed by exsanguination. Through cardiac punctures, terminal sera was collected and stored at -20°C until analysis.

### Competitive sandwich ELISA

Immunoassays were performed as previously described^71^. 384 well ELISA plates (VWR, cat. CA62409-064) were coated with 1 mg/mL NeutrAvidin (Thermo Fisher, cat. A2666) overnight at 4°C and then blocked with 5% BSA for 1 hour at RT. Along with an N-terminus his tag, each TbpB was expressed with a BAP tag, allowing *in vivo* biotinylation upon expression in *E. coli.* Following overnight growth in auto-induction media, as described for dot blots, the lysate was diluted 1:5 in PBS with 1% BSA to wells for 2 hours at RT and then washed with PBS-T thrice. Wells were blocked with 5% BSA for 1 hour at RT again and washed again thrice with PBS-T. Individual mouse serum was diluted (1:1000000) in PBS with 1% BSA and added to their respective wells and incubated overnight at 4°C. The next day, wells were washed with PBS-T thrice and goat α-mouse IgG H&L (HRP) (1:5000, Abcam, cat. ab6789) was added to wells for 2 hours at RT. After a final wash with PBS-T, plates were developed with KPL SureBlue TMB Microwell Peroxidase Substrate (SeraCare, cat. 5120-0077). The reaction was stopped with 2N H_2_SO_4_ and read on a Cytation5 at wavelengths 450 and 570 nm. For hTf blocking, a 1:50 dilution of 1 mg/mL holo-hTf was applied after the incubation with mouse sera. An α-hTf-HRP antibody (Rockland, cat. 600-403-034) was used as a secondary antibody and signal was normalized to a no serum control.

### Optimizing TbpB expression

WHO L was grown overnight on GC agar supplemented with Kellogg’s at 37°C with 5% CO_2_. To determine how to induce TbpB expression, bacteria were grown in two different medias: solid GC Kellogg’s supplemented with 10 µM of deferoxamine mesylate (desferal) or liquid RPMI 1640 (Sigma, cat. R8758). The desferal-supplemented plate was incubated at 37°C with 5% CO_2_, while the bacteria were diluted 1:10 in RPMI 1640 at 37°C for 16 hours with agitation at 160 rpm. Bacteria grown in RPMI were harvested through centrifugation (2000x*g*, 10 minutes). Upon normalization at an OD550 of 0.6, bacteria were boiled at 95°C for 10 minutes in 2X SDS loading dye containing BME. Samples were loaded onto a 12% SDS-PAGE and transferred to a PVDF membrane. The blot was blocked with 5% skim milk in PBS and washed with PBS-T thrice. Antisera collected against 48627^WT^ was used to probe for TbpB expression and goat α-mouse-HRP (1:5000) was employed as a secondary antibody. Finally, membranes were incubated with Enhanced Chemiluminescence substrate at RT in the dark and visualized.

### Serum bactericidal assay

To induce expression of TbpB, which is expressed in low-iron environments, bacteria were collected off the plate with a Dacron swab and resuspended in RPMI 1640 for overnight growth at 37°C shaking at 160 rpm. The next day, bacteria were harvested (3000x*g*, 10 minutes) and resuspended in RPMI 1640. To limit non-specific bactericidal activity, protein A agarose (Thermo, cat. 20333) was incubated with pooled sera acquired prior to challenge. Resin was washed with PBS and eluted with 0.1 M glycine pH 2 (BioShop, cat. GLN002) and buffer exchanged into PBS. Antibodies were filtered (0.22 µm filter, Corning Costar, cat. CLS8161) to remove any aggregates before being stored at -20°C for bactericidal and opsonophagocytic assays. To allow opsonization, approximately 500 bacteria were incubated with antibodies (∼1.25 µg, 0.2 mg/mL) for 20 minutes at 37°C with 5% CO_2_. A mouse monoclonal antibody, 2C7, was provided from Dr. Sanjay Ram’s lab as a positive control (2.5 µg)^36,72^. To initiate the assay, 12% pooled human complement depleted of IgG/IgM (Pel Freez, cat. 34010, Lot 22344) was added and subsequently incubated for 60 minutes at 37°C with 5% CO_2_. Bacteria were plated on GC agar with Kellogg’s at 60 minutes and compared to the no complement control to calculate percent (%) viability.

### Fluorescence microscopy

RAW 264.7 macrophages (ATCC, TIB-71) were cultured in RPMI 1640 with 10% fetal bovine serum (FBS, Gibco, cat. 12483020) to a confluency of over 50%. Before the assay, cells were scraped off and enumerated to seed onto acid-treated glass cover slips at a density of 2*10^4^ cells in RPMI with 10% FBS. Cells were incubated overnight at 37°C with 5% CO_2_ overnight. Iron-starved WHO L was prepared as above and a MOI of 10 was opsonized with antibodies from each immunizing group or the control (RPMI 1640) for 30 minutes at 37°C with 5% CO_2_. Each cover slip was washed gently with pre-warmed RPMI 1640 once before infection with bacteria. Infection was initiated with a 300x*g* centrifugation for 2 minutes at RT. Cover slips were incubated at 37°C with 5% CO_2_ for 1 hour and gently washed with PBS+/+ once with a centrifugation at 500x*g* for 3 minutes at RT. Cells were fixed with 4% paraformaldehyde for 15 minutes at RT and washed with PBS+/+ twice. Cover slips were blocked in 1% BSA for 30 minutes and washed thrice with PBS. Rabbit polyclonal sera raised against *Ngo* MS11 was applied (1:800), followed by goat α-rabbit IgG (H+L) conjugated to Alexa 647 (1:1000 dilution, Invitrogen, cat. A21245). Cells were permeabilized in 0.1% Triton X-100 (octoxynol X-100, BioShop, cat.TRX777.100) in 1% BSA and blocked again in 1% BSA. The same α-gonococcal rabbit polyclonal sera was applied, but differentially visualized with donkey α-rabbit polyclonal conjugated to Alexa 488 (1:1000 dilution, Invitrogen, cat. A11034). Nuclei and cell membrane were visualized with mounting medium with DAPI (Prolong Gold antifade reagent, Life Technologies, cat. P36931) and Phalloidin-Texas Red (1:200 dilution, Invitrogen, cat. T7471) respectively. The Zeiss Observer Z1 epi-fluorescent microscope was used to image each sample. Using at least two representative images per coverslip, thresholds were manually established in Fiji (ImageJ) for the Alexa 647 and Alexa 488 channels, corresponding to extracellular and total bacterial populations, respectively. These thresholds were optimized to identify particles consistent with individual gonococci, enabling quantification within each channel. To estimate the number of bacteria associated with macrophages, the count of extracellular bacteria (Alexa 647) was subtracted from the total bacterial count (Alexa 488) for each image. This difference was then divided by the number of macrophages, quantified manually, to calculate the average number of cell-associated bacteria per macrophage

### Graphing and statistical analysis

Survival curves, immunogenicity graphs, and statistical tests were executed using GraphPad Prism (v. 7.03; GraphPad, San Diego, CA). All data with three or more groups were analyzed using one-way ANOVA with Tukey *post hoc* test for multiple comparisons. Data with two groups were analyzed with unpaired two-tailed Student’s t-test. Colonization curves were analyzed with the Log-rank (Mantel-Cox) statistical test.

## 6. Acknowledgements

This work was supported by a National Institute of Health funding #R01-AI125421-01A1 and #1RO1AI141229. T.F.M. is supported by the Ontario Ministry of Education and Innovation and holds the Canada Research Chair in the Structural Biology of Membrane Proteins, while S.D.G. holds the Canada Research Chair in Infectious Immunopathogenesis. We thank Dr. Zhijie Li, Alan Wong, and Aidan Tomlinson for aiding in structural refinement of MS11^WT-Nlobe^. Diffraction data collection was performed using a beamline CMCF-BM at the Canadian Light Source, a national research facility of the University of Saskatchewan, which is supported by the Canada Foundation for Innovation (CFI), the Natural Sciences and Engineering Research Council (NSERC), the National Research Council (NRC), the Canadian Institutes of Health Research (CIHR), the Government of Saskatchewan, and the University of Saskatchewan. We thank Dr. Muhamed-Kheir Taha for providing the BALB/c expressing hTf mouse line. In addition, we would like to thank Dr. Sanjay Ram who provided the 2C7 monoclonal antibody for validation of serum bactericidal assays, Dr. Charles Deber for providing the CD spectrophotometer and Tracy Stone for help with data collection. Moreover, we are grateful to the animal support staff at the University of Toronto’s Division of Comparative medicine for technical and welfare support of mouse studies performed. We are also thankful to the technical support of Isaac Lee and Dr. Nelly Leung throughout all mouse studies.

